# CD301b lectin expression in the breast tumor microenvironment augments tumor growth

**DOI:** 10.1101/2024.03.13.584829

**Authors:** Ahmet Ozdilek, Amy V. Paschall, Zahra Nawaz, Afaq M. M. Niyas, Fikri Y. Avci

## Abstract

Aberrant tumor glycosylation can alter immune recognition; however, the specific influence of glycan-lectin interactions on tumor progression remains poorly understood. Here, we identify the C-type lectin receptor CD301b (encoded by *Mgl2*) as a regulator of immune activity within the breast tumor microenvironment (TME). Using a murine triple-negative breast cancer model, we demonstrate that tumors expressing the Tn glycoantigen grow more rapidly, and this growth is facilitated by CD301b⁺ immune cells. Depletion or genetic loss of CD301b markedly suppressed tumor growth, indicating that CD301b promotes tumor progression through myeloid-tumor interactions. Phenotypic analyses revealed that CD301b⁺ cells within tumors are type 2 conventional dendritic cells (cDC2s), a subset known to influence immune polarization. Single-cell RNA sequencing of human breast cancers showed that the human ortholog CLEC10A is expressed in cDC2-like dendritic cells and select macrophage subsets, suggesting a conserved role for CD301⁺ myeloid populations. Transcriptomic profiling of tumors developed in *Mgl2*-deficient mice revealed a shift toward an inflammatory, immune-activated state consistent with enhanced antitumor immunity. Together, these findings establish a link between tumor glycosylation and lectin signaling of myeloid cells, highlighting CD301b as a potential target for reprogramming the tumor immune microenvironment in breast cancer.

## Introduction

A major setback observed in many cancers arises from immune modulation within the tumor microenvironment (TME), preventing effective anti-cancer responses. Vaccines, CAR T cells, and checkpoint blockade prime the immune system against cancer-associated antigens, promoting cancer cell destruction through immune cytotoxicity ^1^. Immune therapies for breast cancer remain largely ineffective due to the ability of the tumor to grow unimpeded and eventually metastasize. Immune regulation can occur through cytokine secretion (such as IL-10), signaling mechanisms, or checkpoints (such as through PD-1) ^2–5^. During the malignant transformation of a mammalian cell, a dramatic and aberrant modification of cellular glycosylation is observed. Tumor-associated carbohydrate antigens, or TACAs, can induce immune suppression, allowing cancer cells to evade immune cells ^6–8^. Lectins are carbohydrate-binding proteins that function as receptors for immune cells and can activate immune regulatory pathways through their interactions with TACAs ^9–11^. Similarly, modulating immune profiles in the TME by engaging sialic-acid-binding immunoglobulin-like lectins (Siglecs) is a known TACA mechanism ^12–14^. Thus, elucidating the roles of lectins in regulating the immune response within the TME is becoming increasingly important.

Tn antigen is a TACA ranked as a high-priority cancer-associated antigen based on its antigenicity and oncogenicity ^11,15–19^. Tn is a truncated form of the cell surface O-glycan, consisting of the terminal O-linked N-acetylgalactosamine (GalNAc) attached to serine or threonine. The aberrant glycosylation associated with Tn may occur when the enzyme responsible for O-glycan elongation, T-synthase or its associated chaperone, Cosmc (C1GALT1C1), becomes functionally inhibited ^20,21^. Tn-expressing mucin 1 (MUC1) has been associated with breast cancer cells ^22,23^, especially in triple-negative breast cancer ^24,25^. Modulating MUC1 expression or Tn glycosylation can inhibit tumor growth ^26–28^. However, developing immune responses against Tn-MUC1 has been problematic ^29^.

CD301, also known as macrophage galactose-type lectin (MGL) or CLEC10A, is a C-type lectin receptor (CLR) that binds to Tn antigen on the surface proteins in humans ^19,30^. In mice, the Tn-recognizing homolog of the human CD301 is CD301b, also known as MGL2 ^19,31^. CD301b is primarily expressed by myeloid cells such as dendritic cells (DCs) and macrophages ^17^. Previously, CD301b-expressing myeloid cell populations have been linked with immunosuppressive responses ^16,17,19,32–34^. DCs and macrophages can suppress the proliferation of CD4^+^ effector T lymphocytes through the interaction of MGL with terminal GalNAc residues on CD45 expressed by T cells ^17^. This cell-specific glycosylation of CD45 provided an immunoregulatory pathway, mediated by MGL, thereby controlling effector T cell function. In another study, CD301b^+^ dendritic cells suppressed T follicular helper cells and antibody responses to protein antigens ^32^. Recently, an immunosuppressive DC subset expressing CD301b was shown to accumulate at secondary sites and promote metastasis in pancreatic cancer ^34^ and lung cancer ^35^.

In this study, we investigated the role of CD301b in a murine model of triple-negative breast cancer and found that the loss of CD301b expression significantly restricted tumor growth. Within the tumor microenvironment, CD301b-expressing immune cells were identified as type 2 conventional dendritic cells. Analysis of publicly available single-cell RNA sequencing (scRNA-seq) datasets revealed similar CD301-expressing myeloid cell populations in human breast cancer tissues, suggesting that this regulatory axis may extend beyond the murine model. To further explore the mechanisms underlying the observed tumor growth restriction, we performed bulk RNA sequencing (bulk RNA-seq) on murine tumors, which showed heightened inflammatory immune responses when CD301b was absent. Together, these findings link CD301 expression to the control of tumor-associated inflammation and point to its potential as a target for new breast cancer therapies.

## Results

### CD301b/Tn Axis Impacts Tumor Growth

We first aimed to investigate the relationship between CD301b^+^ immune cells and Tn glycoantigen-expressing breast cancer cells. We used a CRISPR-Cas9 gene editing model to knock out Cosmc expression in AT3 murine breast cancer cells. Cosmc is a chaperone essential in elongating the core O-glycan beyond the truncated Tn form of α-GalNAc ^20,21,36^. When the Cosmc function/expression is disrupted, elongation of the O-glycan is not observed; instead, the Tn antigen is observed at significantly higher levels. After disrupting Cosmc expression in these cell lines, we confirmed increased Tn cell surface expression through flow cytometry using both a reBaGs6 IgM antibody (Suppl. Fig. 1A) ^37^ as well as a complementary biotinylated VVL lectin coupled with fluorescent streptavidin (Suppl. Fig. 1B). Both staining methods indicated significantly higher Tn expression on the *Cosmc* KO cell line. We also isolated RNA from each cell line and confirmed decreased Cosmc expression in the Tn^hi^ cell line through qPCR using *Cosmc* primers (Suppl. Fig. 1C). We then tested whether knocking out Cosmc expression changes the proliferation rate of the AT3 breast cancer cells *in vitro*. After culturing both cell lines at the same concentrations for three days, we observed no significant differences in cell proliferation between the two lines (Suppl. Fig. 1D), indicating that Tn expression alone does not promote cancer cell growth.

To determine *in vivo* tumor cell growth, we injected AT3 (Tn^lo^) and AT3 *Cosmc* KO (Tn^hi^) murine breast tumor cells into the mammary pads of C57BL/6 mice and monitored tumor growth. We observed that Tn^hi^ tumors grew significantly faster than Tn^lo^ tumors (Fig. 1A), indicating that Tn expression impacts tumor growth rate.

**Fig. 1.**
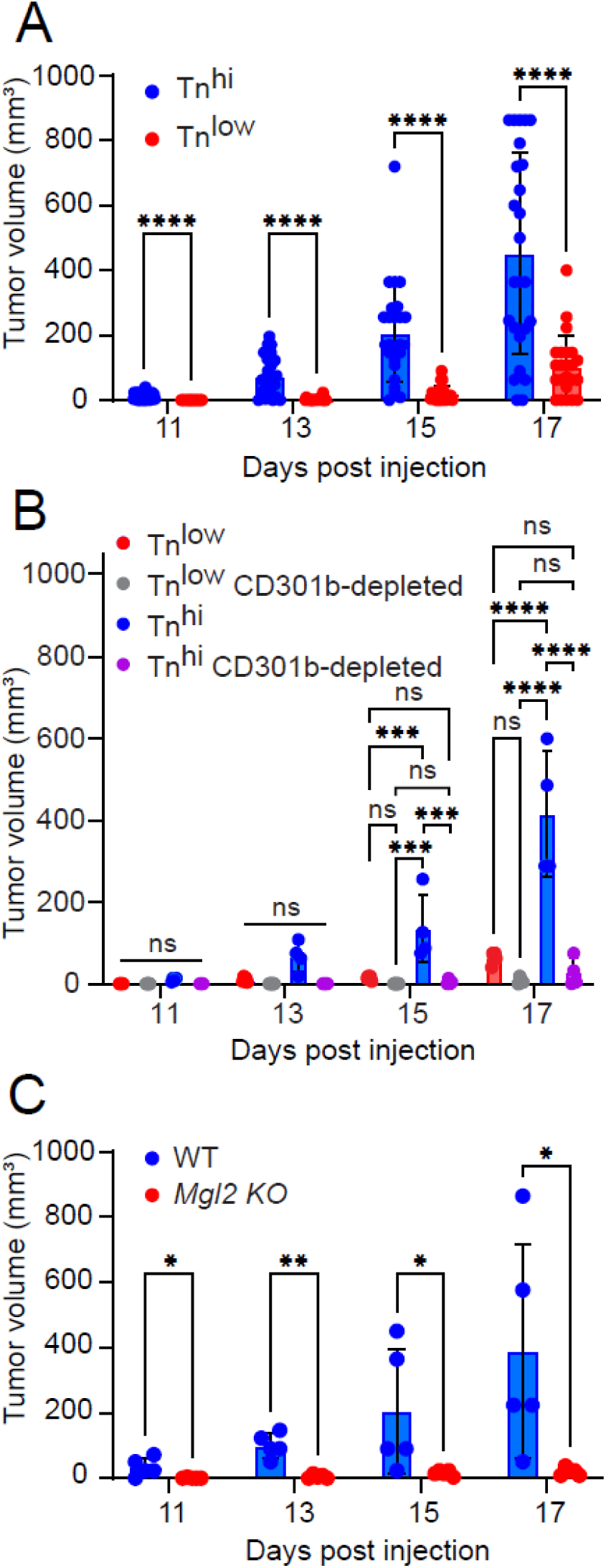
CD301b/Tn Axis Impacts Tumor Growth. **A.** AT3 (Tn^low^) and AT3 *Cosmc* KO cells (Tn^hi^) (2.5E5/mouse) were injected into the mammary pads of C57BL/6 mice (n = 25). Tumor sizes were monitored. Tumor sizes were calculated based on (length times width^2^)/2 for volumes in mm^3^. **B.** AT3 (Tn^low^) and AT3 *Cosmc* KO cells (Tn^hi^) tumor cells (2.5×10^5^/mouse) were injected into the mammary pads of WT mice and heterozygous Mgl2-DTR mice (n= 4 or 5) with or without CD301b^+^ cells depleted. Tumor sizes were monitored. **C.** AT3 *Cosmc* KO cells (Tn^hi^) (2.5×10^5^/mouse) were injected into the mammary pads of *Mgl2* KO mice (n=5). Tumor sizes were monitored.

To examine the contribution of CD301b-expressing immune cells to tumor growth, we employed heterozygous *Mgl2^+/DTReGFP^* mice (Mgl2-DTR), in which CD301b+ immune cells can be selectively depleted by diphtheria toxin (DT) treatment ^32,33^. In the first experiment (Fig. 1B), AT3 Tn^low^ or Tn^hi^ tumor cells were injected into Mgl2-DTR mice with or without DT administration. Tumor growth was significantly reduced in DT-treated mice of the Tn^hi^ group, indicating that CD301b^+^ immune cells promote tumor progression. In a complementary experiment (Fig. 1C), we injected AT3 Tn^hi^ cells into homozygous *Mgl2^DTReGFP/DTReGFP^* mice (CD301b-null, *Mgl2* KO), which lack surface expression of CD301b due to the disruption of both alleles but retain the immune cell populations. These mice also exhibited markedly reduced tumor growth compared with wild-type controls. Together, these experiments indicate that the observed phenotype is associated with tumor Tn expression and facilitated by the CD301b protein.

### Tumor-infiltrating CD301b^+^ cells display a type 2 conventional dendritic cell (cDC2) phenotype

We next characterized tumor-infiltrating CD45^+^CD301b^+^ immune cells in the murine triple-negative breast cancer model. These cells expressed CD11c, a canonical dendritic cell (DC) marker ^38,39^ (Fig. 2A) and were strongly positive for MHCII, confirming their DC identity (Fig. 2B). CD301b⁺ cells also expressed CD11b, a defining marker of mouse type 2 conventional dendritic cells (cDC2s) ^40^ (Fig. 2B). Mouse cDC2s can be distinguished from cDC1s by CD103 and SIRP-alpha expressions ^38,41^. We examined the expression of these markers by tumor-infiltrating CD301b^+^ cells, which display a cDC2s phenotype (Fig. 2C). Although CD301b^+^ cells are primarily cDC2s, not all DCs or cDC2s express CD301b in the TME (Fig. 2D and 2E).

**Fig. 2.**
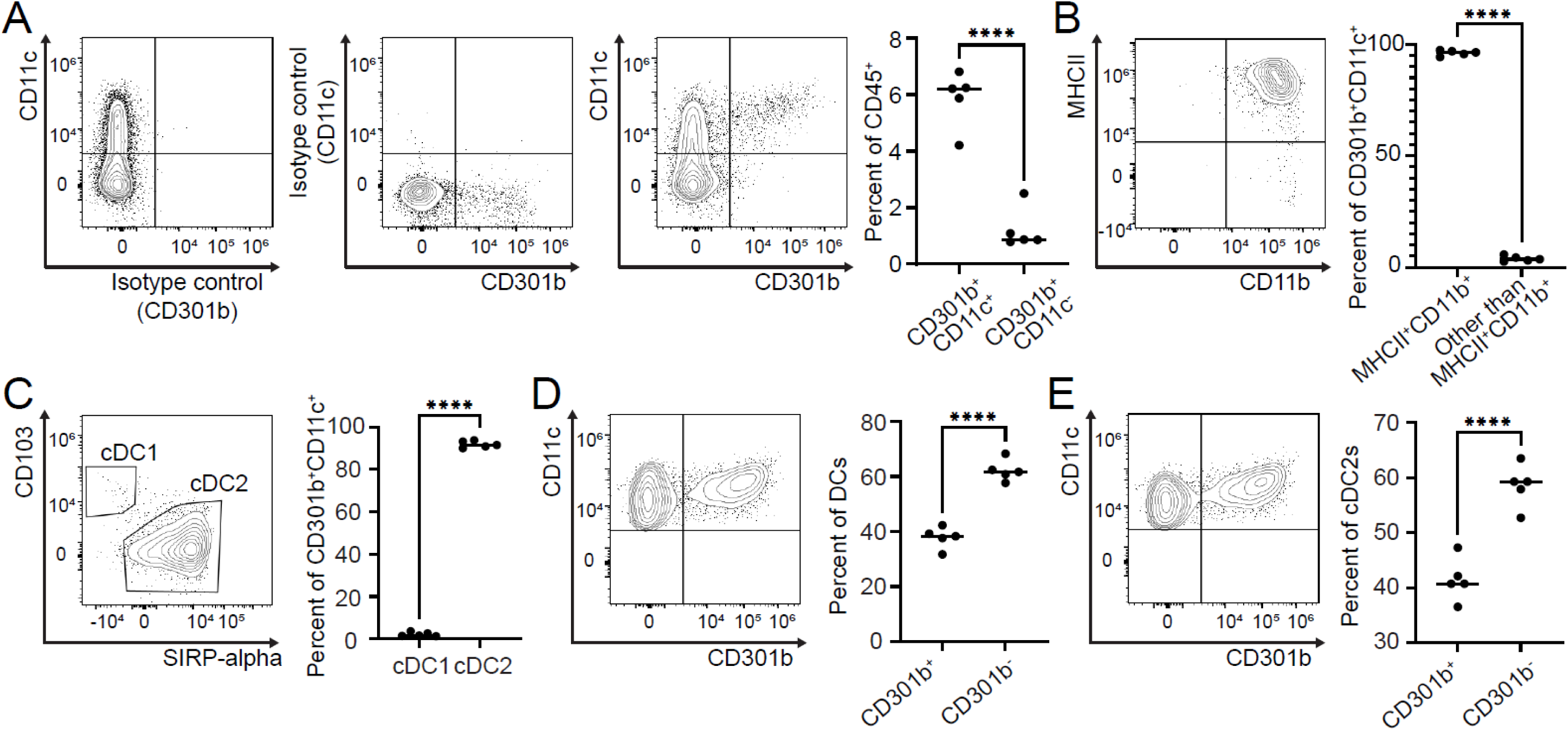
Characterization of CD45^+^CD301b^+^ immune cells in murine breast cancer TME. Single cell suspensions from tumors of WT mice injected with AT3 *Cosmc* KO cells (Tn^hi^) were stained, and expressions of surface markers were analyzed with flow cytometry (gated as in Suppl. Fig. 2). **A.** Among live CD45^+^ cells, CD301b^+^ cells are CD11c^+^. **B.** CD301b^+^ and CD11c^+^ cells are MHCII^+^ and CD11b^+^. **C.** CD301b^+^ and CD11c^+^ cells are cDC2. DCs and cDC2s in the TME, **D and E**, respectively, consist of Cd301b-negative and positive cells.

cDC2s constitute a subset of antigen-presenting cells that play key roles in coordinating adaptive immune responses. In contrast to cDC1s, which specialize in cross-presentation and cytotoxic T cell activation, cDC2s promote CD4⁺ T cell priming and modulate immune polarization within tissues ^42–44^. Recent studies have shown that cDC2s exhibit remarkable plasticity in the tumor microenvironment, where they can adopt either immunostimulatory or tolerogenic phenotypes in response to local cues ^44^.

### CD301⁺ immune cells in the human breast cancer TME include both dendritic cells and macrophages

Since CD301b⁺ cells in the murine tumor microenvironment were identified as cDC2s, we then investigated whether CD301 expression in human breast cancers similarly correlated with dendritic cells or extended to other myeloid subsets. To address this, we analyzed publicly available single-cell RNA sequencing (scRNA-seq) data (GSE161529) from 20 patients, including triple-negative (n = 8), ER⁺ (n = 6), and HER2⁺ (n = 6) tumors ^45^. Following quality control, data integration and annotation, we focused on CD45⁺ immune cells to map CLEC10A (human CD301) expression across myeloid populations (Fig. 3A; Suppl. Fig. 3A). CLEC10A expression was most prominent in dendritic cells and was also detectable in macrophage/monocyte populations, with negligible expression in other immune cells (Fig. 3B–C). Across breast cancer subtypes, dendritic cells consistently showed higher CLEC10A expression than macrophages (Suppl. Fig. 3B). Within the dendritic cell compartment, we identified four subsets—cDC1, cDC2, cDC-LAMP3⁺, and plasmacytoid DC (pDC)—and found that CLEC10A expression was highest in cDC2 (56.9%) and moderate in cDC-LAMP3⁺ (11.3%), but low in cDC1 (1.9%) and absent in pDCs (0%) (Fig. 3D–F; Suppl. Fig. 3C–D).

**Fig. 3.**
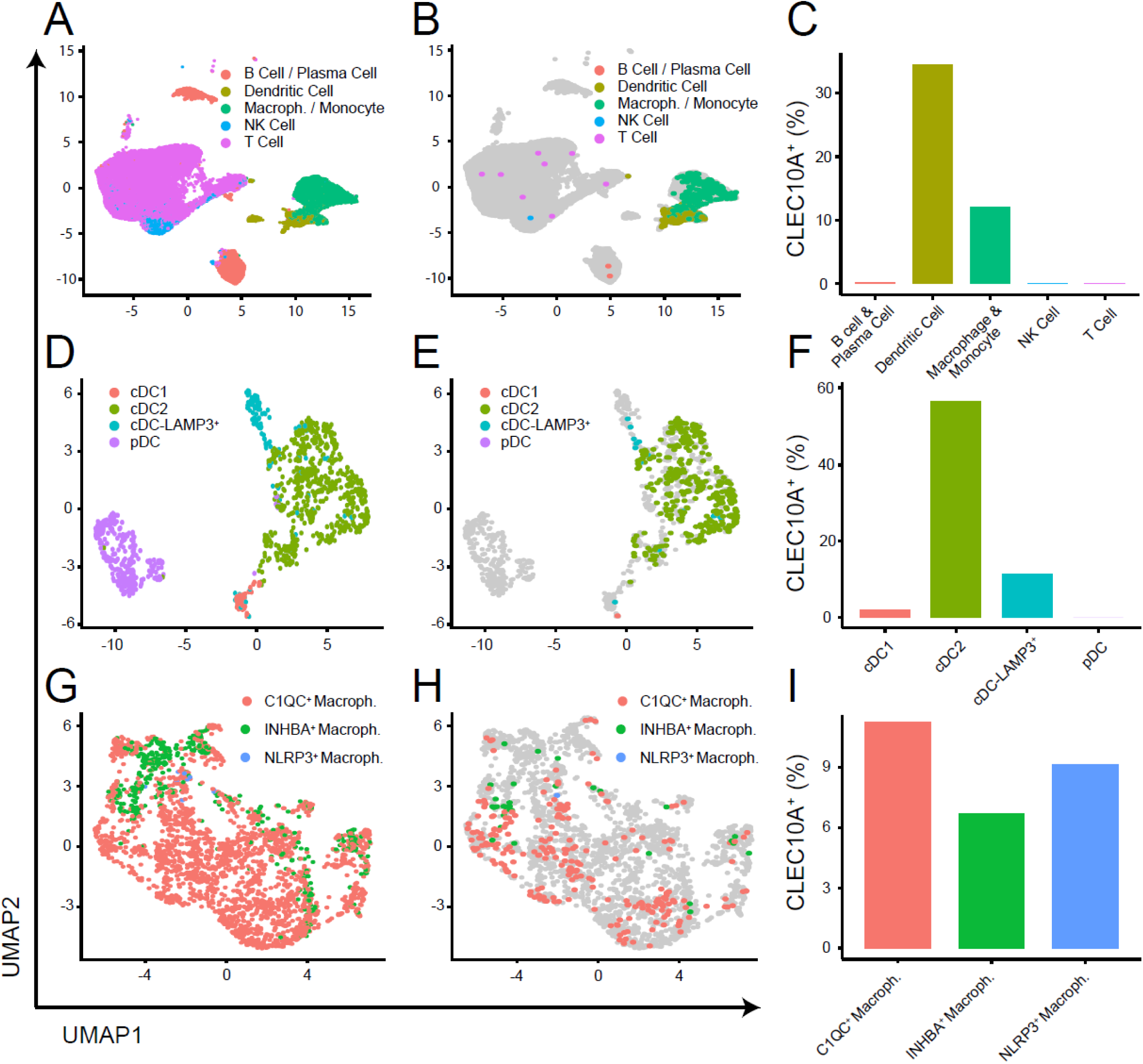
Characterization of CD45^+^CD301b^+^ immune cells in human breast cancer TME. **A.** UMAP of all immune cells colored by major lineage (B/Plasma cell, Dendritic cell, Macrophage/Monocyte, NK cell, T cell). **B.** Immune cells highlighting CLEC10A⁺ cells (colored) over all cells (gray). **C.** Bar plot showing the fraction of CLEC10A⁺ cells within each major lineage. **D.** UMAP of dendritic cell compartment colored by subset (cDC1, cDC2, cDC-LAMP3^+^, pDC). **E.** Dendritic cell subset highlighting CLEC10A⁺ cells (colored) over all DCs (gray). **F.** Bar plot showing the fraction of CLEC10A⁺ cells within each DC subset. **G.** UMAP of macrophage compartment colored by subset: C1QC⁺ macrophages, INHBA⁺ macrophages, and NLRP3⁺ macrophages. **H.** Macrophage subset highlighting CLEC10A⁺ cells (colored) over all macrophages (gray). **I.** Bar plot showing the fraction of CLEC10A⁺ cells within each macrophage subset. *UMAP axes indicate the first two dimensions. Percentages in bar plots are calculated as (CLEC10A⁺ cells / total cells) within the indicated group*.

We next examined tumor-associated macrophages (TAMs). TAMs are key regulators of tumor inflammation, tissue remodeling, and immune suppression ^46^. TAMs were subdivided into transcriptionally defined subsets reflecting distinct functional programs: C1QC⁺ macrophages, associated with immunosuppression ^47–49^; NLRP3⁺ macrophages, associated with poor prognosis and tumor growth ^50,51^; and INHBA⁺ macrophages, linked to angiogenesis, matrix remodeling, and tumor progression ^52^ (Fig. 3G; Suppl. Fig. 3E). Among these subsets, *CLEC10A* transcription distributed similarly in C1QC⁺ macrophages (11%), NLRP3⁺ macrophages (9%), and INHBA⁺ macrophages (7%) (Fig. 3H–I). Interestingly, CLEC10A-positive NLRP3⁺ macrophages were detected exclusively in triple-negative breast cancers, where expression levels were comparable to those in INHBA⁺ macrophages (Suppl. Fig. 3F). This enrichment suggests that CD301 expression extends beyond dendritic cells to select macrophage populations, particularly those engaged in inflammatory and tissue-remodeling responses within aggressive tumor subtypes.

Taken together, these results indicate that CD301 expression in the human breast cancer microenvironment is concentrated within myeloid lineages—encompassing both cDC2-like dendritic cells and specialized macrophage subsets. This pattern mirrors the cellular distribution observed in the mouse TME, suggesting that CD301 marks a conserved myeloid program potentially involved in coordinating immune regulation and tissue remodeling during tumor progression.

### The lack of CD301b is associated with a strong inflammatory immune signature in the breast TME

Since CD301b directly associates with tumor growth and identifies cDC2 and macrophage populations within the breast tumor microenvironment (TME), we then examined how its loss impacts immune signaling and tumor–immune interactions. To this end, we performed bulk RNA sequencing (bulk RNA-seq) on tumors derived from wild-type (WT) and *Mgl2* knockout (*Mgl2* KO) mice following injection of AT3 Tn^hi^ tumor cells. The global transcriptomic heatmap (Fig. 4A) revealed distinct clustering and clear separation between WT and *Mgl2* KO mouse tumors, indicating a strong transcriptional divergence associated with *Mgl2* loss. This observation provided the foundation for downstream pathway and gene-level analyses to elucidate how CD301b influences immune regulation within the TME.

**Fig. 4.**
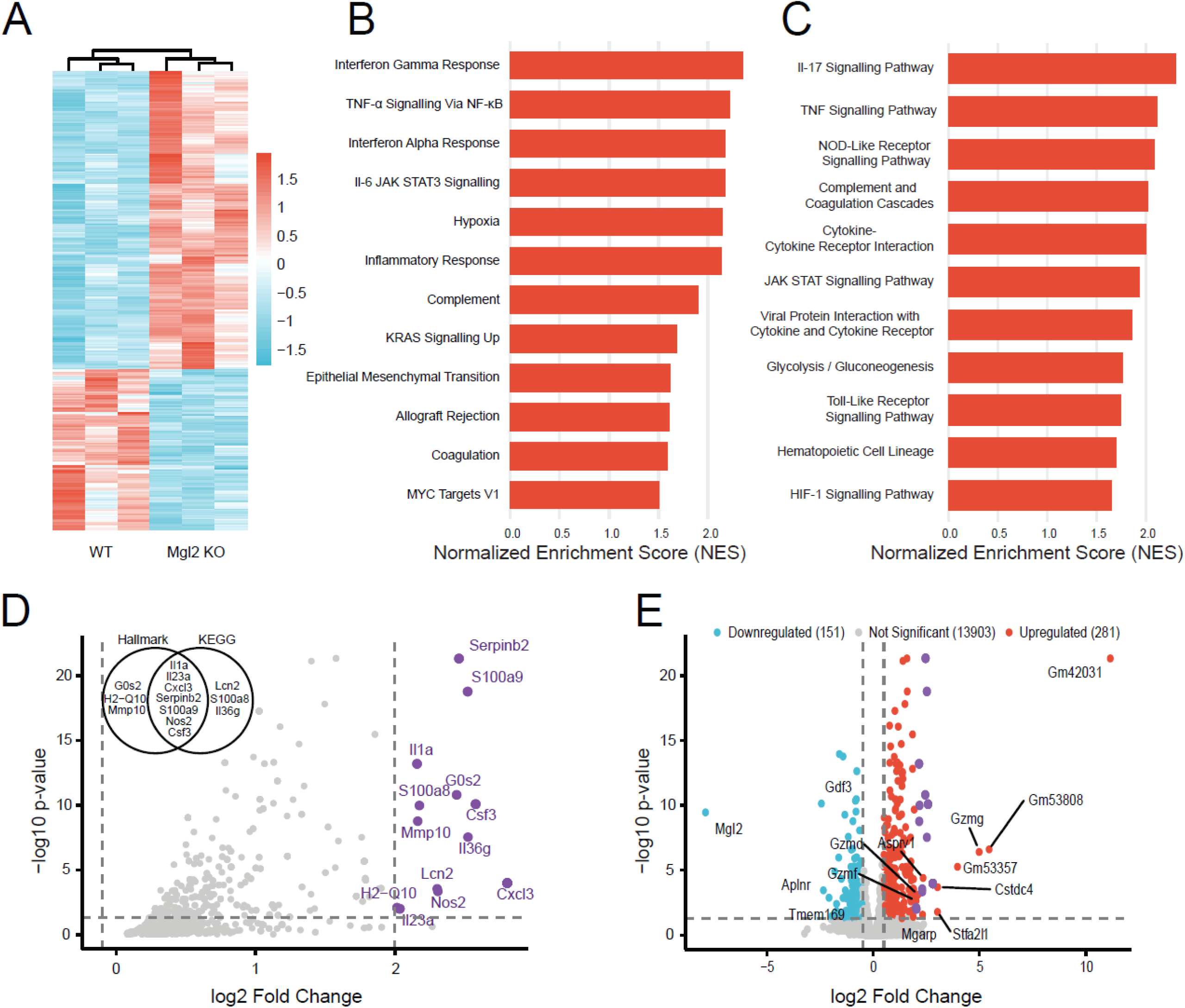
The lack of CD301b is associated with a strong inflammatory immune signature in the breast TME. **A.** Heatmap showing differentially expressed genes (DEGs) identified by RNA-seq from tumors in *Mgl2* KO and WT mice. The blue and red bands indicate low and high gene expression quantity, respectively. Biological replicates showed the highest degree of correlation. **B.** Gene Set Enrichment Analysis (GSEA) of Hallmark pathways reveals immune-related and cancer-associated pathways, all of which are upregulated in Mgl2 KO. **C.** GSEA of KEGG pathways shows immune-related and cancer-associated pathways, and they are all upregulated in Mgl2 KO. **D.** Volcano plot showing the pathway-associated genes (from panels B and C). The genes with a log₂ fold change of 2 or greater are highlighted in purple. The X-axis represents log₂-transformed fold change, and the Y-axis represents −log₁₀-transformed significance. The VENN diagram shows the distribution of genes between Hallmark and KEGG gene sets. **E.** Volcano plot of DEGs between tumors from *Mgl2* KO and WT mice. Red points indicate upregulated DEGs, blue points indicate downregulated DEGs, gray points represent non-significant genes, and purple points indicate the pathway-associated genes (from panel D). All the genes with a log₂ fold change ≥ 2 are labeled.

To dissect these transcriptomic differences, Gene Set Enrichment Analysis (GSEA) was performed using the Hallmark and Kyoto Encyclopedia of Genes and Genomes (KEGG) enrichment gene set databases (Fig. 4B–C). *Mgl2* KO tumors displayed significant enrichment of immune and inflammatory pathways, including TNF-α/NF-κB, IL-6/JAK-STAT3, interferon-α/γ, and IL-17 signaling—pathways broadly associated with myeloid activation, cytokine production, and tumor immunosurveillance ^53,54^. Despite the slower tumor growth observed in the knockout mice, complement/coagulation, hypoxia, and glycolytic pathways were also upregulated, suggesting a compensatory increase in metabolic and inflammatory activity within the TME. Collectively, these data indicate that CD301b functions as an immunoregulatory node that tempers cytokine and interferon responses, whereas its loss enhances pro-inflammatory signaling. The dominance of interferon- and NF-κB–driven programs aligns with the slower tumor progression observed in KO mice, implying that CD301b deficiency reprograms the TME toward a functionally immune-active, tumor-controlling state.

The pathway-focused volcano plot (Fig. 4D) highlights key upregulated genes underpinning these responses, including *Cxcl3, Il1a, Il23a, Il36g, Csf3, Nos2, S100a8, S100a9,* and *Lcn2*. These represent canonical NF-κB and IL-17 targets known to mediate myeloid recruitment, nitric oxide production, and acute-phase inflammation—hallmarks of innate immune activation ^55–57^. The upregulation of the serine protease inhibitor and matrix metalloproteinase genes *Serpinb2* and *Mmp10* in the tumor microenvironment of knockout mice suggests a tumor-suppressive function through the regulation of extracellular matrix remodeling and modulation of immune responses ^58,59^. Thus, CD301b loss produces a highly activated, cytokine-rich TME that maintains elements of tissue repair while fostering effector cell response and immune surveillance.

The volcano plot of all differentially expressed genes (DEGs) (Fig. 4E) contextualizes these changes within the full transcriptome. Upregulated genes largely mirrored those driving enriched pathways, confirming that *Cxcl3, Il1a, Il23a, Nos2, S100a8/a9,* and *Csf3* constitute the core *Mgl2* KO transcriptional program rather than isolated pathway artifacts. On the other hand, downregulation of the TGF-β family growth differentiation factor 3 and the apelin receptor genes *Gdf3* and *Aplnr* is associated with promoting tumor growth, angiogenesis, and metastasis ^60,61^.

Together, these results suggest that CD301b deficiency reprograms the TME toward a cytokine-driven, inflammatory, and interferon-dominant state that enhances immune activation and tumor control. The loss of CD301b lifts an immunoregulatory brake within the TME, unleashing broad inflammatory signaling that, despite introducing metabolic stress, creates a net anti-tumor environment characterized by immune activation and delayed tumor growth.

## Discussion

Aberrant glycosylation is a defining hallmark of malignant transformation, and our findings identify CD301b as a key immunoregulatory lectin linking breast tumor-associated Tn antigens to myeloid immune modulation. CD301b⁺ cells, primarily cDC2s, promoted breast tumor growth, whereas their depletion or genetic loss limited progression. These observations align with previous studies demonstrating that TACAs interact with lectins to influence myeloid differentiation and immune regulation ^19,21,22,62–64^.

Transcriptomic profiling of tumors developed in *Mgl2*-deficient mice revealed broad activation of NF-κB, IL-6–JAK–STAT3, and interferon pathways, consistent with a shift toward a pro-inflammatory, “immune-hot” microenvironment ^65^. These data position CD301b as an immunoregulatory node that tempers innate activation, similar in concept to a checkpoint-like mechanism operating within the myeloid compartment ^9,10,66,67^. While this study does not define the signaling circuitry involved, it indicates that CD301b expression in cDC2s and macrophages contributes to a previously unrecognized immunoregulatory phenotype in breast cancer TME.

The complex and pleiotropic nature of CD301b’s activity may explain its influence on the immune landscape of cancer ^66–68^. CD301b’s modulatory behavior mirrors that of other C-type lectin receptors ^69^. CD301b-mediated restraint may protect against chronic inflammation, but in tumors, it can inadvertently favor immune escape ^70,71^. Conversely, CD301b loss triggers inflammation driven by NF-κB and interferons, which enhances effector recruitment but imposes metabolic and hypoxic stress on the TME ^72^. Together, these findings support a model in which CD301b functions as a molecular regulator balancing immune activation and tolerance in breast cancer.

Mechanistically, CD301b–Tn interactions may regulate antigen processing, cytokine release, or costimulatory signaling within dendritic cells, thereby shaping T cell activation thresholds ^69^. These possibilities remain speculative but highlight the potential of CD301b as a new immune-regulatory axis distinct from canonical checkpoints, such as PD-1 or CTLA-4. Future work should investigate how TACA ligands and intracellular adaptors regulate CD301b signaling and consequent immune programming in the TME. Further single-cell and spatial analyses of breast TME transcriptome in the presence and absence of CD301b will help define how CD301b⁺ subsets integrate into existing immune networks across tumor stages.

From a translational perspective, targeting the CD301–Tn interaction offers a promising route to remodel the breast cancer immune microenvironment. Pharmacologic blockade or glycomimetic interference could complement checkpoint inhibitors by dismantling glycan-mediated myeloid suppression, whereas selective induction of this pathway may have therapeutic relevance in autoimmune disease ^73^. Ultimately, elucidating the intricate mechanisms that govern immune modulation through CD301b will be essential to selectively induce these properties in disease-specific contexts, enabling the development of knowledge-based, precision immunotherapies.

In summary, this study identifies an immunomodulatory CD301b⁺ myeloid phenotype that contributes to breast cancer growth and whose loss induces a robust inflammatory program in the breast TME. While the molecular mechanisms remain to be defined, our data indicate that CD301b acts as a glycan-sensitive, checkpoint-like regulator of myeloid activity. Elucidating its ligand specificity and downstream signaling will clarify how breast tumor glycosylation reshapes TME and may reveal new strategies to enhance immunotherapy efficacy.

## Materials and Methods

### Mice

Eight-week-old female C57BL/6 mice were obtained from Jackson Laboratories (Bar Harbor, ME) and housed at Emory University Whitehead Biomedical Research Building. *Mgl2^DTReGFP/DTReGFP^* mice were a generous gift from Akiko Iwasaki at Yale University. To obtain heterozygous *Mgl2^+/DTReGFP^* mice, C57BL/6 were bred with *Mgl2^DTReGFP/DTReGFP^*mice. Mice were kept in microisolator cages and handled under biosafety level 2 (BSL2) hoods. For tissue processing and subsequent flow cytometry, mice were euthanized by carbon dioxide inhalation in accordance with IACUC guidelines. Where applicable, cell suspensions were generated through mechanical tissue disruption and collagenase D digestion. Red blood cells were lysed, and samples were filtered through 60 μm nylon filters to obtain single-cell suspensions. For depletion of CD301b^+^ cells, heterozygous Mgl2-DTR mice were treated with diphtheria toxin (0.5μg/mouse) in sterile PBS intraperitoneally every two to three days, starting at day -1 before tumor injection.

All mouse experiments were in compliance with the Emory University Institutional Animal Care and Use Committee under an approved animal use protocol. Our animal use protocol adheres to the principles outlined in *U.S. Government Principles for the Utilization and Care of Vertebrate Animals Used in Testing*, *Research and Training*, the Animal Welfare Act, the *Guide for the Care and Use of Laboratory Animals*, and the *AVMA Guidelines for the Euthanasia of Animals*.

### Generation of Tn^hi^ breast cancer cells

To express Tn glycans at high levels in tumor cells, we used a CRISPR/Cas9 methodology to stably silence Cosmc expression in AT3 cells using established protocols and reagents. Mouse *Cosmc* guide RNA and CRISPR/Cas9 plasmid were obtained from Santa Cruz Technology (sc-425587). AT3 murine breast cancer cells were a generous gift from the Kebin Liu lab at Augusta University. AT3 cells were transfected with *Cosmc* CRISPR/Cas9 KO plasmid according to the manufacturer’s protocol. Puromycin was used to select transfected cells. We then used flow cytometry to confirm higher expression of Tn on the AT3 cell surfaces using the ReBaG6 antibody (generously provided by Richard Cummings at Harvard University) (Suppl. Fig. 1A) ^37^ and VVL lectin (Vector Laboratories) (Suppl. Fig. 1B). The proliferations of transfected and untransfected cell lines (AT3 Tn^hi^ and AT3 Tn^low^) were tested in an MTT proliferation assay for three days, and colorimetric analysis was performed with a CytoTek plate reader according to protocol; no significant differences in proliferation compared to parent cells were observed (Suppl. Fig. 1D). Cell lines were maintained in RPMI media supplemented with 10% FBS, sodium pyruvate, HEPES buffer, NEAA, β-mercaptoethanol, and penicillin/streptomycin at 37°C, 5% CO2.

### AT3 Tn^low^ and/or AT3 Tn^hi^ Tumor Challenge

AT3 cells were harvested and washed in sterile PBS. Cells were suspended in a final concentration of 2.5E6/ml sterile PBS. Cells were subcutaneously injected into the mammary pads of mice at 2.5E5/100μl/mouse. Mice were observed and euthanized at the tumor endpoint (maximum tumor dimension between 0.9 cm and 1.2 cm. Tumor volumes were calculated as ((length x width x width)/2) in mm^3^.

### Flow Cytometry

Cells were stained in PBS with TruStain fcX (BioLegend, Cat. No. 101320) to reduce non-specific antibody binding. Cell samples were stained with the following antibodies and stains: CD11c-PacBlue (BioLegend, clone N418), CD11b-Alexa Fluor 488 (BioLegend, clone M1/70), SIRPα-Alexa Fluor 700 (BioLegend, clone P84), CD103-PE/Dazzle 594 (BioLegend, clone 2E7), MHCII-BV785 (BioLegend, clone M5/114.15.2), CD45 Alexa Fluor 647 (BioLegend, clone 30-F11), CD301b-PE (BioLegend, clone URA-1), and LIVE/DEAD Fixable Blue (Invitrogen). All isotype controls were obtained from BioLegend. Samples were washed and analyzed with flow cytometry (Cytek Aurora). Fluorescence minus one (FMO) plus specific isotype control antibody-stained samples were used as negative staining controls, and single stains were used for compensation. Flow cytometry data were analyzed using FlowJo Single Cell Analysis Software with gating strategies shown in Suppl. Fig. 2.

### Bulk RNA-seq and Bioinformatics

Tumors were harvested from wild-type C57BL/6J and *Mgl2* KO mice (n = 3 per group). Total RNA was extracted using the MagMAX™ mirVana™ Total RNA Isolation Kit in combination with the KingFisher Apex system (Thermo Fisher Scientific). RNA integrity and concentration were assessed with a Qubit 3.0 fluorometer (Thermo Fisher Scientific). High-quality RNA samples were used for library preparation, followed by quality control and sequencing using Novogene’s standard protocol. Libraries were sequenced on the Illumina NovaSeq X Plus platform to generate paired-end 150 bp reads (PE150) at Novogene Inc. Raw FASTQ files were retrieved and subjected to quality control with FastQC ^74^. Reads were aligned to the *Mus musculus* reference genome (GRCm39/mm39) using HISAT2 ^75^. A gene-level count matrix was generated with featureCounts ^76^. The count matrix was imported to the downstream differential expression analysis using the *DESeq2* R package ^77^. Significantly differentially expressed genes were defined by adjusted p-value < 0.05. Gene Set Enrichment Analysis (GSEA) was performed on the RNA-seq dataset using the clusterProfiler R package ^78^ to identify significantly enriched KEGG and hallmark pathways between *Mgl2* KO and wild-type tumors. To focus on cancer-relevant biology, enrichment results were refined to include immune- and cancer-related pathways.

### Analyzing the scRNA-seq data

The scRNA-seq data generated using the 10x Genomics Chromium platform from human breast cancer tumors (GSE161529) ^45^ were obtained from Gene Expression Omnibus (GEO) ^79^. Data were processed into the Seurat R package (version 5), and low-quality cells were removed based on established Seurat quality control parameters ^80^. After QC, putative doublets were removed using the *DoubletFinder* R package ^81^. Datasets were integrated, and non-immune cells were removed by subsetting the CD45⁺ cells with detectable *PTPRC* expression (*PTPRC* > 0). Following preprocessing, cell annotation was contacted with scATOMIC ^82^. The subclasses of the dendritic cells and macrophages were further validated with canonical lineage markers ^83,84^.

## Statistical Analysis

GraphPad Prism v8 was used for statistical analyses. Two-way ANOVA with Tukey’s multiple comparisons test was used to determine statistical significance between experimental groups in each of the applicable experimental models (Fig. 1B). An unpaired parametric two-tailed t-test was used for Fig. 1A, 1C, 2A, 2B, 2C, 2D, 2E, and Suppl. Fig. 1C. Significance is indicated on each graph based on p-value: >0.05 = ns; <.05 = *; <0.01 = **; <0.001 = ***; <0.0001 = ****.

## Acknowledgements

We acknowledge the following individuals for their contributions of transgenic mice, cell lines, and monoclonal antibodies: Dr. Akiko Iwasaki, Dr. Kebin Liu, and Dr. Richard Cummings. National Institutes of Health grants R01AI123383 and R01AI152766 supported this work.

## Notes

### Competing Interest Statement

The authors have declared no competing interest.

### Summary of Updates

In this revised manuscript, we have made several updates to improve clarity, accuracy, and completeness. We have added two new authors, one of whom is now listed as the primary author to reflect their central contributions to the new analyses and manuscript development. We have also incorporated two new transcriptomic datasets. The first is a human breast cancer single-cell RNA-seq dataset, and the second is a mouse breast tumor microenvironment bulk RNA-seq dataset. These datasets are now fully described in the Methods section, and the corresponding analyses have been integrated into the Results and figures. Their inclusion provides additional context and strengthens the overall interpretability of the study. Additionally, we repeated all flow-cytometry analyses presented in the original submission to ensure consistency and robustness. Through these updated analyses, we more clearly characterized the CD301b⁺ immune cell populations examined in the study. Based on phenotypic markers and updated gating strategies, we now identify these CD301b⁺ cells as a subset of dendritic cells. The revised data enable us to better define these CD301b-expressing cells within the broader immune compartment.

