## Supplementary figures and images for "CD301b lectin expression in the breast tumor microenvironment augments tumor growth"

### Supplemental Figures

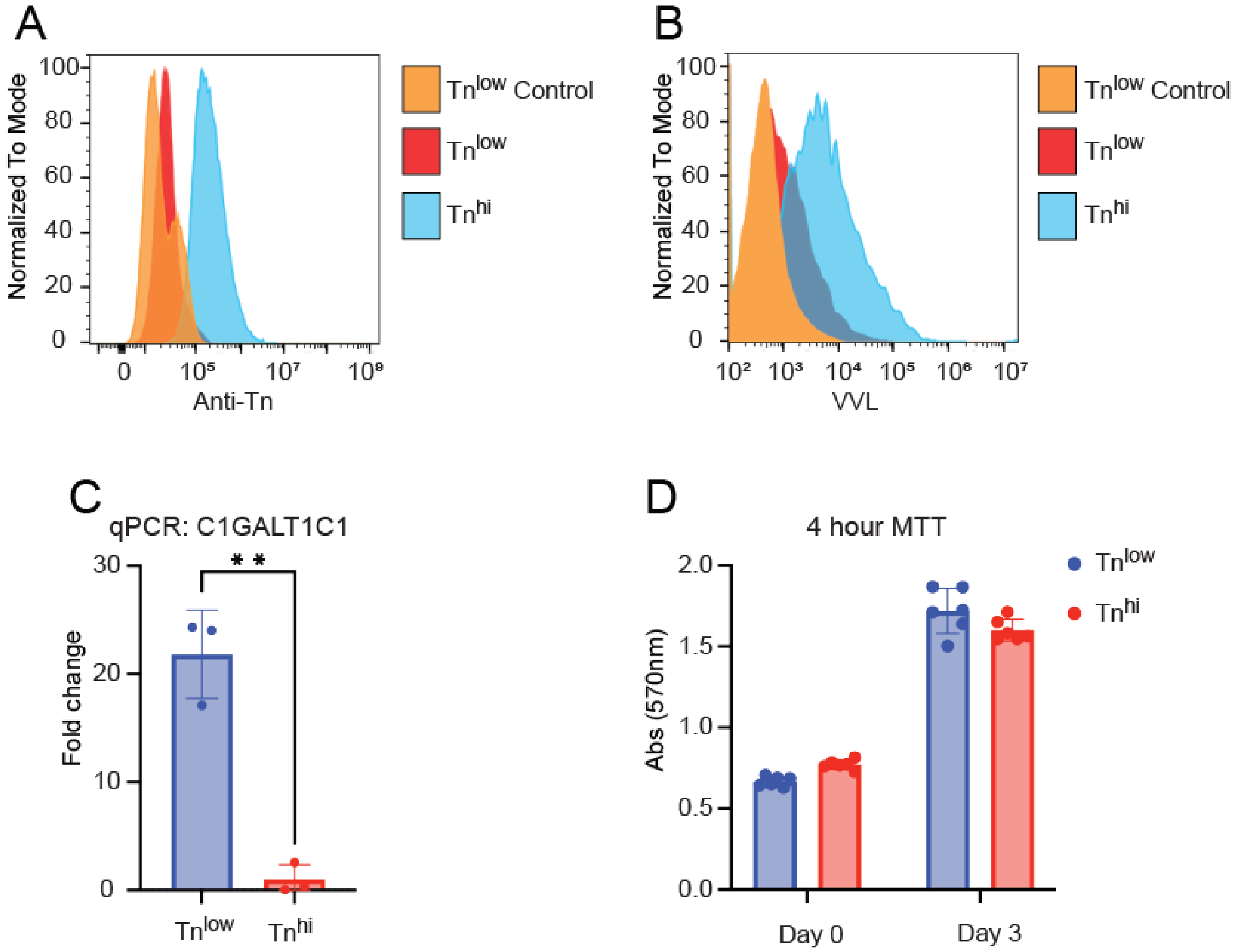

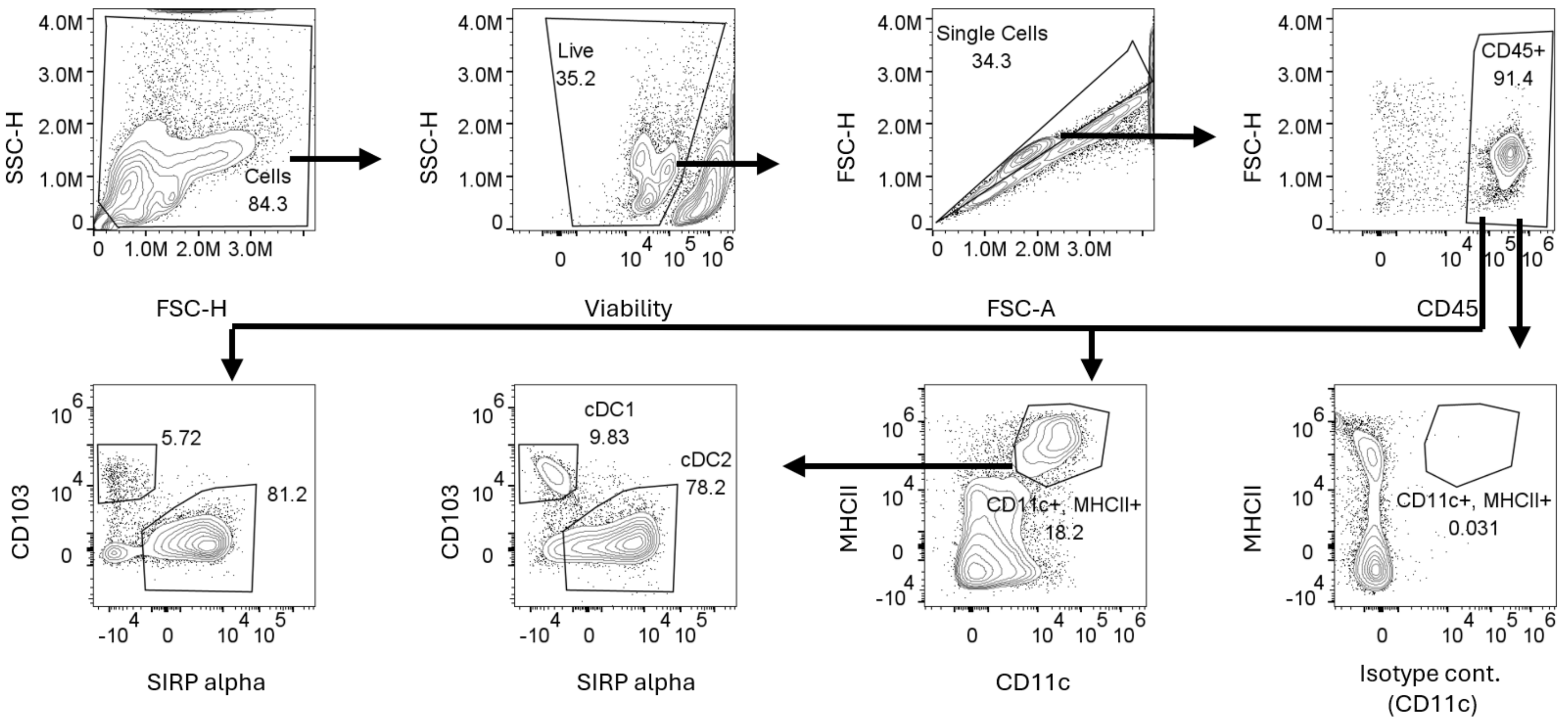

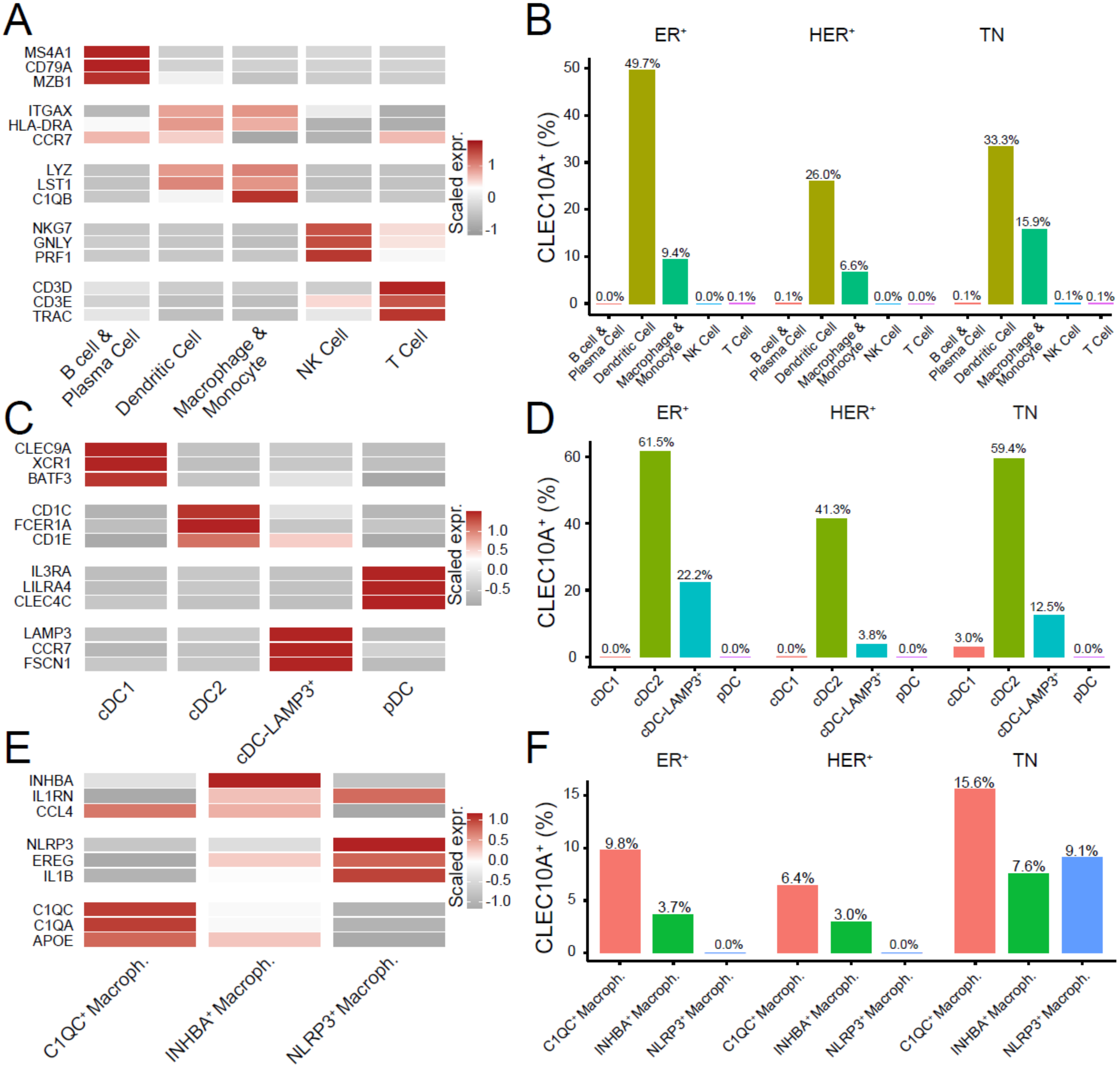
